# ACCURATE DECODING OF THE SPINAL CORD OUTPUT IN HUMANS WITH IMPLANTED HIGH-DENSITY ELECTRODE ARRAYS

**DOI:** 10.1101/2022.01.29.478247

**Authors:** Silvia Muceli, Wigand Poppendieck, Aleš Holobar, Simon Gandevia, David Liebetanz, Dario Farina

## Abstract

Invasive electromyography opened a new window to explore motoneuron behaviour *in vivo*. However, the technique is limited by the small fraction of active motoneurons that can be concurrently detected, precluding a population analysis in natural tasks. Here, we developed a high-density intramuscular electrode for *in vivo* human recordings along with a fully automatic methodology that could detect the discharges of action potentials of up to 67 concurrently active motoneurons with 99% accuracy. These data revealed that motoneurons of the same pool receive common synaptic input at frequencies up to 75 Hz and that late recruited motoneurons inhibit the discharges of those recruited earlier. These results constitute an important step in the population coding analysis of the human motor system *in vivo*.

## INTRODUCTION

The introduction of intramuscular needles and wires for electromyography (EMG) by Adrian and Bronk (1929) and Basmajian and Stecko (1962) opened a window to explore the neural underpinning of movement control. By recording muscle fibre action potentials, intramuscular EMG reveals the timing of the action potentials discharged by the innervating spinal motoneurons (MN). The analysis of motor units (MUs) from intramuscular EMG decomposition rapidly became the standard approach to study MN behaviour *in vivo* in humans and other species (Desmedt, 1973).

Nonetheless, the use of EMG to assess MNs also imposes some constraints. Some intramuscular electrodes are highly selective to detect the electrical activity of a small number of muscle fibres. This makes it easy to identify the discharge times of a few MUs through EMG decomposition, which is conventionally based on spike sorting of action potentials with similar morphology (LeFever and De Luca, 1982). However, the electrode selectivity implies that only a small fraction of the hundreds of active MNs can be studied concurrently. To increase the number of sampled MUs, investigators have serially recorded single MU activity. While serial recordings have unravelled patterns of MN firing, a MN population analysis is still missing, which limits our understanding of the process of generation of the neural output of the spinal cord. Currently, there is no robust method that provides simultaneous decoding of a large portion of the active MNs in natural tasks.

The identification of large populations of concurrently active MUs is necessary to characterise the synaptic inputs received by MNs. Coherence among spike trains of the homonymous MN pool reflects the common synaptic input at various frequency bands. A single MN cannot accurately sample an input with a frequency greater than half its average discharge rate (Lazar and Pnevmatikakis, 2008; Lazar and Tóth, 2004), which is usually in the range 10 - 40 Hz (Enoka and Fuglevand, 2001). As a result, sampling by few MNs limits the frequency range at which coherence (and thus common synaptic input) can be observed. However, as the common synaptic input is spread to the whole MN pool (Farina et al., 2014), pooling the spike trains extracted from large populations of MUs allows sampling at higher frequencies.

As a further example, analysis of the output of a population of MNs is also a way to investigate connectivity among MNs, e.g. due to Renshaw inhibition (Eccles et al., 1961; Renshaw, 1941). Renshaw cells receive collateral projections from MN axons and synapse on MNs mediating recurrent inhibition back to the MN pool. However, the distribution of recurrent inhibition throughout the MN pool is unknown in humans (Alvarez, 2019). Most knowledge about recurrent inhibition stems from experiments on anesthetized animal preparations, and direct translation of findings to human studies of intact MNs during natural behaviour is challenging. Again, technological advances for sampling large populations of MUs *in vivo* in humans are necessary (Alvarez, 2019).

A way to increase the number of concurrently detected MUs in natural tasks uses decomposition of activity recorded with high-density grids of surface electrodes (Holobar et al., 2009). However, surface EMG only detects the activity of superficial MUs (Farina et al., 2010). As an alternative approach to increase the number of sampled MUs, we previously introduced multichannel intramuscular electrodes based on thin-film technology (Farina et al., 2008; Muceli et al., 2015), which provide a large and unbiased sample of MUs from both deep and superficial muscles. These electrodes comprise a linear array of detection points in a flexible wire that can record across the muscle cross-section. Tens of MUs can be concurrently detected with these systems (Muceli et al., 2015). Yet, these systems are limited to only 16 electrode sites and they require partially manual spike sorting. Spike sorting software for multichannel intramuscular EMG indeed currently relies on human oversight to edit the results (McGill et al., 2005).

When increasing the number of recorded signals, the EMG decomposition process must be applied to each recorded EMG channel. With conventional spike sorting, this increases computation time as well as manual editing of the results (Enoka, 2019). Alternative to spike sorting, blind source separation (BSS) methods can be applied to separate sources (MUs) when a large number of observations (EMG channels) is available (Negro et al., 2016). However, classic BSS limits the maximum number of extracted sources to the number of observations (in practice to less than the observations).

Here, we describe two breakthroughs in the technology to investigate MN behaviour *in vivo*. First, we designed, manufactured and tested a novel implantable electrode array for human studies with a much greater number of recording sites and higher site density than any previous systems. The novel design allowed the implantation of the array acutely with needles of similar size to those used in conventional concentric needle recording. Second, we used a fully automatic decomposition algorithm (no manual editing) that enabled the decoding of the high-density multiunit recordings with accuracy comparable to that achieved by extensive manual editing of each trace by an expert operator. Further, with this new technology, we addressed two fundamental open questions in MN physiology. We found that a MN pool receives common synaptic input is a frequency range up to 75 Hz, much greater than previously thought. We then analysed the effect of individual MU discharges on the MN population output to determine the connectivity among MNs.

## RESULTS

### Intramuscular thin-film electrode array

We designed and manufactured a high-density intramuscular array with 40 platinum electrodes of area 5257 μm^2^ each (Fig. 1 A), linearly distributed over a 2-cm length. Figure 1B shows the complete layout of the double-sided thin-film structure. The structure is built on a polyimide substrate, has a total length of 7 cm and is U-shaped with two filaments of width 655 μm and 150 μm (Fig. 1C), and thickness of 20 μm. The wider filament contains two linear arrays of 20 oval electrodes each (Fig. 1A), with 1-mm inter-electrode distance on the top (cyan) and bottom (green) sides of the polyimide (Fig. 1C). The two arrays have a shift of 0.5 mm (Fig. 1C). Since the double-sided structure is only 20-μm thick, it is equivalent to a linear array of electrodes with 0.5 mm inter-site distance. The number of electrodes is limited by the number of interconnection lines fitting on the filament. The advantage of two arrays on the two sides of the structure is that the filament width can be reduced for a given number of electrodes. Also, the occurrence of short-circuits during manufacturing is reduced. The narrower filament is inserted into a 25-gauge needle (100 Sterican, B. Braun, Melsungen, Germany), to introduce the thin-film structure into a muscle, with a procedure similar to that used in classic fine wire EMG. The needle is withdrawn leaving the array inside the muscle.

**FIGURE 1:**
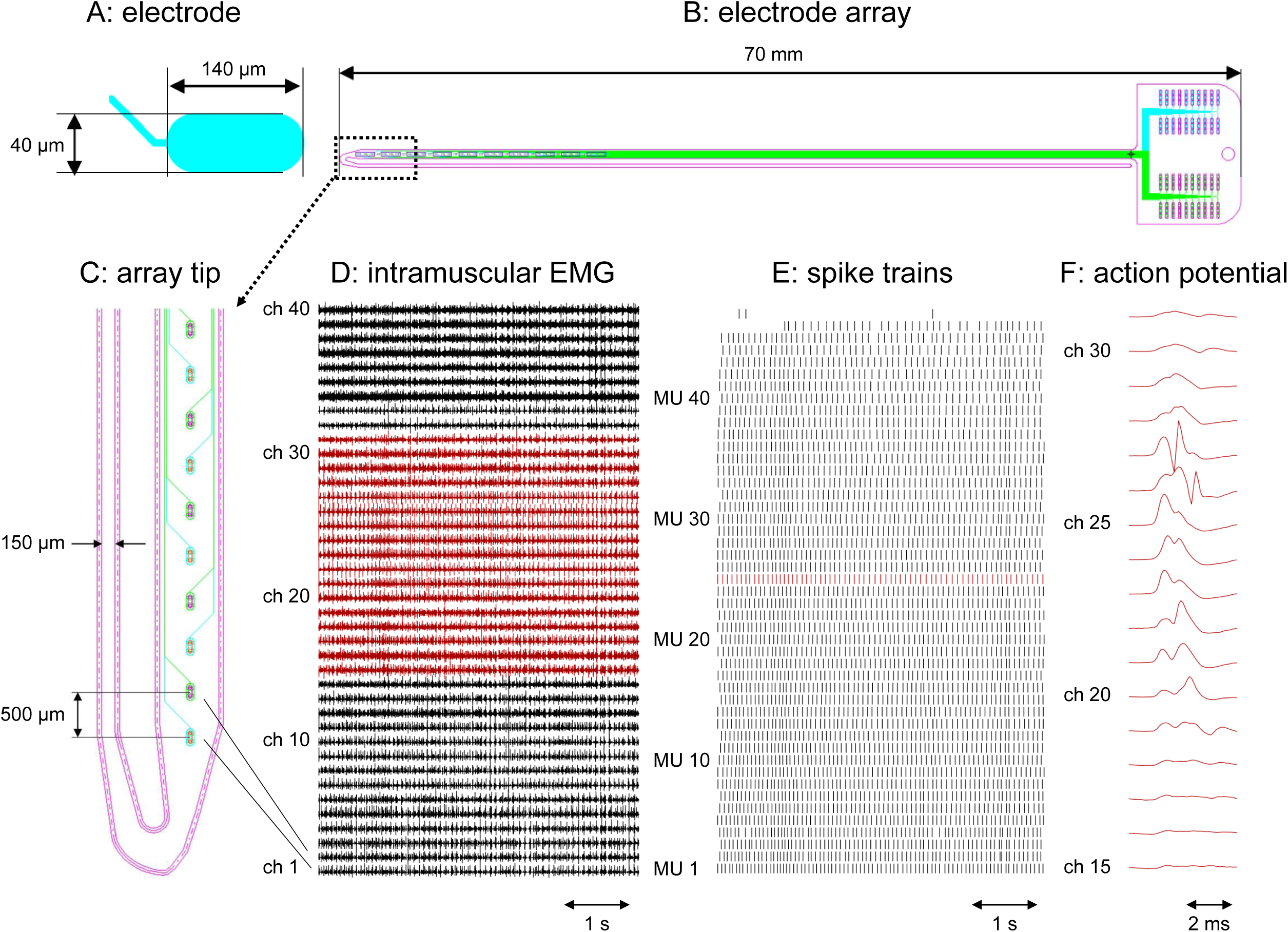
Design of the double-sided electrode array and representative recordings. (A) Close-up of an oval electrode. (B) Whole structures with the tracks running towards the connection pad. (C) Close-up of the electrode array tip. Electrodes represented in cyan are located on the top side of the thin-film array and those in green are located on the bottom side of the wider filament. (D) Representative recordings obtained from the tibialis anterior of S1 during a contraction at 20% of the maximal force (MVC). (E) Firing pattern of 45 MUs extracted from the signal shown in D. (F) Multichannel action potentials of a representative motor unit obtained by averaging the red-coloured EMG channels in panel D with the firing pattern of the same colour in panel E as a trigger.

### Signal quality and motor unit yield

The electrode array was tested in three healthy men (S1-S3). Two arrays were inserted in the tibialis anterior of subject S1, while one array was implanted in the other two subjects. S1 performed a steady contraction at 20% of maximal force (MVC), whereas S2 and S3 contracted the tibialis at 30% MVC. The electrodes recorded high quality signals, with a baseline noise of 15.8 ± 9.9 μV (average ± standard deviation across 4 arrays of 40 channels each). Figure 1D displays representative signals recorded from S1 to show the signal-to-noise ratio. Figure 1E shows the firing patterns of the MUs extracted via manual decomposition from the signals recorded from array1 in S1. In the raster plot, each row represents a different MU, and each vertical line the discharge of an action potential. Within the selected time frame (5 s), 45 MUs were consistently active, 1 MU was recruited during the contraction and 1 had a few isolated discharges. Figure 3F shows a representative example of a MU action potential detected across several electrodes of the 40-channel array.

The recorded signals were decomposed independently into the constituent MU action potential trains by two expert investigators (SM and AH). We refer to the two decomposition processes as manual and automatic decomposition. For manual decomposition, intramuscular EMG signals from each thin-film system were decomposed channel by channel using spike sorting software (McGill et al., 2005), manually edited for resolving missed discharges and superimpositions, and merged (after resolving differences in the discharge patterns of the same MU extracted from different channels) so that each MU’s activity was represented by a unique firing pattern. For automatic decomposition, all signals from the same array were decomposed with the BSS method (see Methods and Holobar and Zazula (2007)). We then compared the MU firing patterns extracted by the two decomposition procedures (manual and automatic) via the rate of agreement (RoA).

Table 1 reports the data obtained via the decomposition process. The activity of 161 MUs was manually decomposed from the signals recorded from the 4 arrays, yielding 38735 unique discharges in 20 s. The RoA between all possible pairs of MUs detected from the same array (1225, 630, 741, 630, for S1 array1, S1 array2, S2, and S3, respectively) ranged from 0 to 11%, confirming that all identified MU spike trains had few common discharges, i.e., they were unique. The number of channels in which the peak-to-peak amplitude of the corresponding action potential exceeded 10 times the RMS baseline noise ranged from 4 to 40 (median 18) for all MUs but 3 (148 MUs in total). The presence of the same MU over multiple channels contributed to the accurate extraction of the MU firing patterns (Mambrito and De Luca, 1984). The average firing rate was 14.8 ± 1.7, 11.0 ± 1.2, and 12.7 ± 1.9 Hz for S1-3, in agreement with previous studies (Connelly et al., 1999; Erim et al., 1996). Most MUs were active for the whole 20 s interval, but 10 of 161 fired less than 50 times each and were excluded from the calculation of the average firing rate and number of channels exceeding baseline to increase the reliability of the estimates. There were no MUs in common between array1 and array2 of S1 (RoA between all possible pairs (1800) ranged between 0 and 5%). The cross-spike triggered averaging procedures produced averages at the baseline noise level, further confirming that there were no MUs in common between array1 and array2.

**TABLE 1:**
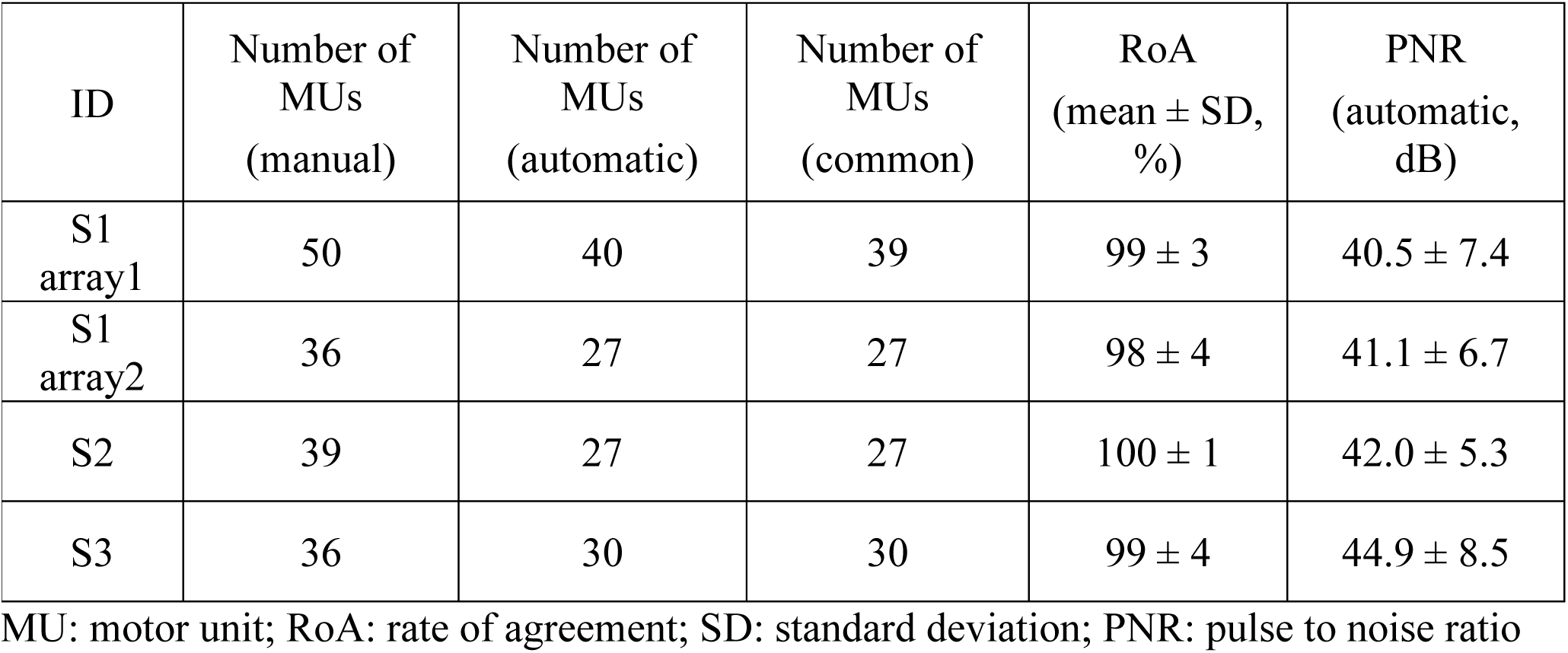
Decomposition performance for the high-density intramuscular signals: manual versus automatic decomposition

| ID | Number of MUs (manual) | Number of MUs (automatic) | Number of MUs (common) | RoA (mean ± SD, %) | PNR (automatic, dB) |
| --- | --- | --- | --- | --- | --- |
| S1 array1 | 50 | 40 | 39 | $99 \pm 3$ | $40.5 \pm 7.4$ |
| S1 array2 | 36 | 27 | 27 | $98 \pm 4$ | $41.1 \pm 6.7$ |
| S2 | 39 | 27 | 27 | $100 \pm 1$ | $42.0 \pm 5.3$ |
| S3 | 36 | 30 | 30 | $99 \pm 4$ | $44.9 \pm 8.5$ |

MU: motor unit; RoA: rate of agreement; SD: standard deviation; PNR: pulse to noise ratio

### Decomposition accuracy

The manual decomposition of each channel (20-s recording) took >8h by the expert operator. The fully automatic decomposition of each array (40 channels, 22 s) took 2h and 9 min of computational time on average across the 4 arrays (Intel CORE i9 vPro 9Gen Processor with 32 GB RAM). Table 1 includes the comparison between the output of the manual and automatic decomposition procedures. About 80% of the MUs identified by manual decomposition were identified by the automatic decomposition. Only one MU identified by automatic BSS did not match a MU extracted by manual decomposition. The investigator who performed the manual decomposition initially identified the unmatched MU, but she discarded it from further analysis because of lack of confidence in the decomposition accuracy due to the low amplitude of its action potentials. Eight MUs that were not extracted by the automatic decomposition (21%) fired less than 50 times.

The average RoA across the 123 MU spike trains that were identified by both procedures (manual and automatic) was 99 ± 3%. Of those 123 spike trains, 64 matched the automatic results with a 100% RoA, and 36 had a RoA ≥ 99%. We inspected the disagreement between the output of the two procedures and found that only 3 common MUs had a RoA in the range 80 to 85% due to misalignments in discharge timings which was greater than our strict threshold of 0.5 ms. One of the three MUs had a satellite action potential. Among the common MUs, 16 discharges identified by the manual decomposition and missed by the automatic decomposition were doublets.

Taken together, these results indicate that the high-density intramuscular array yields high MU sampling and the activity of most of the MUs can be reliably extracted by a fully automatic procedure with comparable accuracy to manual decomposition.

### Motor unit population coherence

We calculated the coherence between groups of MUs of increasing numerosity (Fig. 2). Figure 2A shows 20 s of spike trains extracted from S1. Figure 2B shows the corresponding coherence for groups of MUs between 1 and 34. The coherence was statistically significant (i.e., above the 95% confidence level) for frequencies of about 75 Hz, proving that the synaptic input bandwidth goes well beyond the β band. Similarly, the coherence was still significant at ∼75 Hz for S3 (Fig. 2D). In both cases, an increase of coherence in the gamma band with the number of MUs is clear. On the contrary, for S2, the coherence bandwidth was limited to 40 Hz (Fig. 2C).

**FIGURE 2.**
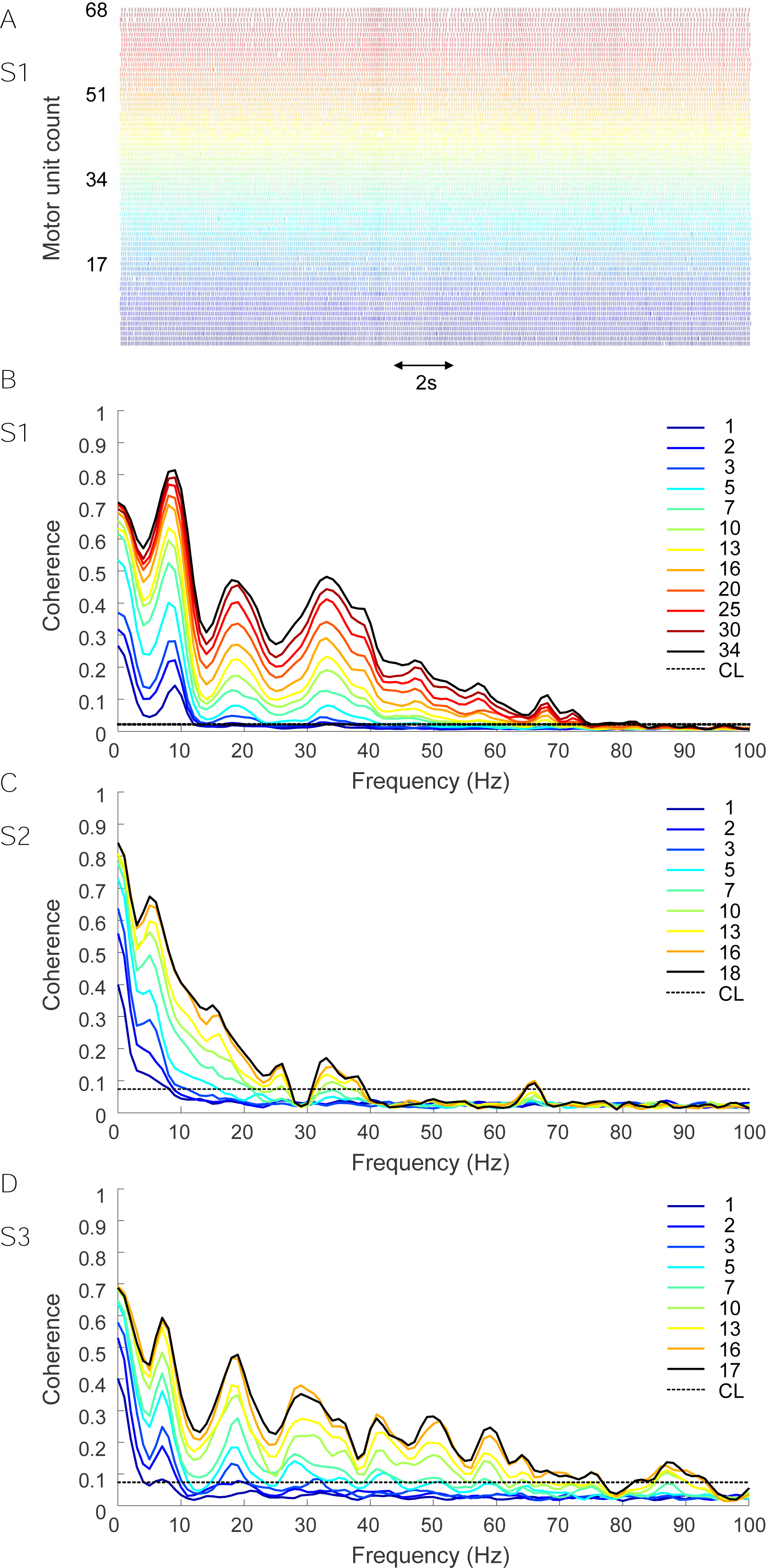
Coherence between populations of motor units. (A) Firing pattern of 68 motor units active during 20 MVC contraction (S2, 2 arrays). Coherence between combinations of cumulative spike trains (CSTs) obtained by pooling an increasing number of motor units from subject S1 (B), S2 (C), and S3 (D). Black dashed horizontal line is the 95% confidence limit. Coherence increased with the motor unit numerosity and the population coherence was significant up to 40 Hz in S2, and up to 75 Hz in S1 and S3, respectively. Note: 60 s of data were used for S1, 20 s for S2 and S3.

### Reciprocal effect of motoneuron discharges on the homonymous pool

The discharge of a MN depends on supraspinal and spinal inputs, including from interneurons. A particular class of interneurons, the Renshaw cells, cause recurrent inhibition of the homonymous MN pool (Hultborn et al., 1979). Renshaw cells are facilitated during weak and inhibited during strong contractions (Hultborn and Pierrot-Deseilligny, 1979). We expected to see the effects of reciprocal inhibition in our recordings when the subject exerted forces of 20 or 30% MVC. As there are opposing views on the distribution of recurrent inhibition between early- and late-recruited MUs within the same MN pool (Granit et al., 1957; Haase et al., 1975; Hultborn et al., 1988), we separately investigated higher and lower threshold MUs. Results are reported in Fig. 3 as synchronization cross-histograms. Firing rate was considered a surrogate of recruitment order, in that early recruited MUs discharge faster, at a given moderate level of force, than those recruited later (De Luca and Erim, 1994). As can be observed in both S1 and S3, late recruited MNs caused more inhibition of the discharges of the early recruited MNs at ∼15 ms (dip in Fig. 3 A and C) than the converse. On the other hand, for S2 (Fig. 3B), inhibition continued up to ∼40 ms. No dips were observed in the cross-histograms obtained by applying different perturbations (see METHODS, Connectivity among motoneurons) to the original firing patterns and maintaining the firing rate unchanged (control condition; results not shown), implying that the latter did not influence the results presented.

**FIGURE 3.**
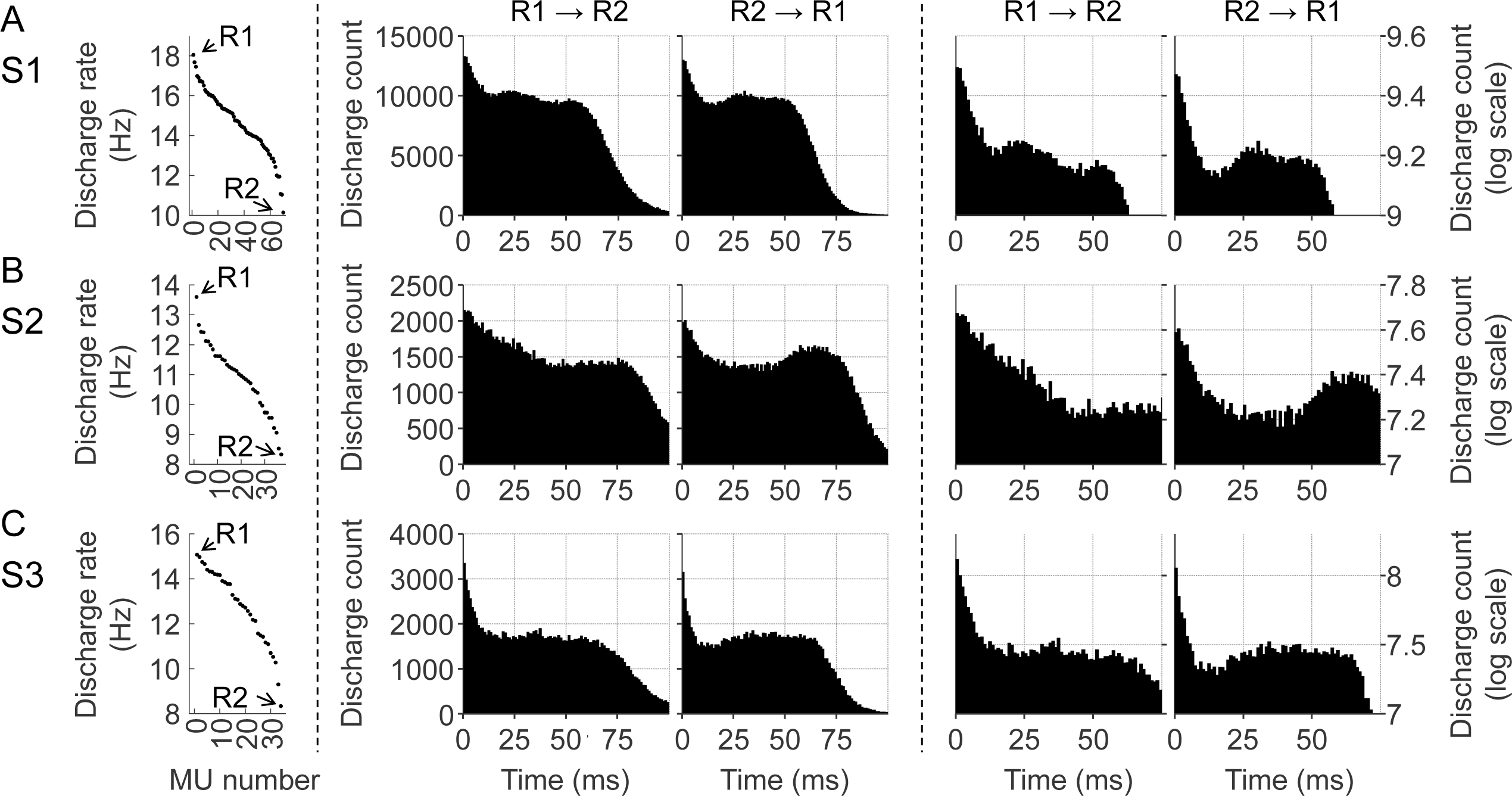
Analysis of motor unit synchronization for subjects S1 (A), S2 (B), and S3 (C). Left panels show the average discharge rate of the motor units in a 20 s time interval. Central R1→ R2 (R2 → R1) panels display the influence of earlier (later) recruited motor units on the discharge timing of the later (earlier) recruited motor units via cross-histograms between pairs of motor unit spike trains. The two rightmost columns represent the same values in logarithmic scale so that the inhibition can be more readily visualised.

## DISCUSSION

We have presented the development of a high-density electrode array for intramuscular recordings that enables the automatic accurate extraction of tens of MUs concurrently active. We have shown representative examples of MU population analysis enabled by our system.

### Intramuscular array

Our electrode array configuration consists of polymer (Hassler et al., 2011) and metal that are micromachined (Stieglitz et al., 2000) into a thread containing 40 electrodes. The materials and minimal thickness (20 μm) confer the required flexibility to interface the muscle without being unpleasant for the subject. Each electrode has an area of 5257 μm^2^. Such small electrodes inevitably present high electrical impedance which reduces the signal-to-noise ratio. The contacts were therefore coated with microrough platinum that increases the active surface and reduces the impedance by 10 times compared to an untreated electrode (Muceli et al., 2015, 2019). The array has electrodes manufactured on both sides of the substrate (Poppendieck et al., 2015) to enable increased spatial resolution and to reduce the likelihood of short-circuits. This improvement in the technology allowed us to build 40 electrodes in a 2-cm long filament.

### Motor unit decomposition

Four intramuscular electrode arrays were tested in 3 subjects. Electrodes were inserted into the tibialis anterior and used to acquire EMG during isometric contractions at moderate force. Each array yielded an average number of 40 concurrently active MUs. Eighty-six MUs could be extracted from a contraction at 20% MVC with two high-density electrode arrays in S1. Given that the tibialis anterior is assumed to comprise about 450 MUs (Enoka, 1995) and the relatively low muscle force exerted by S1, the identified 86 MUs represent a relative large proportion of those that were active during the contraction.

On average, 31 MUs per array could be automatically decomposed with an accuracy of 99% when compared with manual expert decomposition. Compared to previous systems with fewer electrodes (Muceli et al., 2015), the number of automatically extracted MUs with the proposed high-density electrode is 2 to 3 times greater and the accuracy substantially higher (Negro et al., 2016). For example, our previous attempt at automatic decomposition of EMG recorded with two arrays of 16 channels each yielded 22 out of 53, 24 of out 57, and 21 out of 60 (i.e., about 40%) manually detected MUs at different force levels, with an average RoA of 94%. Our high-density system enabled automatic decomposition of about 80% of the manually detected MUs action potential trains constituting the interference EMG with a 99% RoA. Eight MUs identified by manual decomposition discharged less than 50 times, which was insufficient for the automatic identification. The yield of MUs per channel was also superior to that achieved by BSS of high-density surface EMG data from the tibialis anterior (21 MUs/64 channels) (Del Vecchio et al., 2020) that in any case can only detect MUs with large action potentials at the skin surface.

The automatic decomposition was validated against the manually decomposed dataset. The RoA between the two procedures was 99% on average (across 123 MUs). This value is remarkably high and can be attributed to the high-density of channels. The comparison between the two decomposition procedures is a conservative approach for estimating accuracy. As signals were decomposed independently by two decomposition methods and operators, the likelihood that the same mistake is made in the two cases is very low (Mambrito and De Luca, 1984). Therefore, the procedure of validation of the automatic decomposition in this study is robust. In addition, the average pulse-to-noise ratio across the 124 MUs automatically extracted was 42 dB, greater than values reported for surface EMG decomposition (Holobar et al., 2014), further confirming the high accuracy of the automatic decomposition procedure.

We inspected the disagreement between the two decomposition procedures, and we identified two sources of errors (doublets and misalignments). Some of the doublets could not be identified by the automatic BSS decomposition. This is to be expected as doublets may have an action potential with smaller amplitude compared to the main action potential when the second input (forming the doublet) arrives at the end-plate before the muscle had fully recovered (Denslow, 1948). As the BSS algorithm can only identify action potentials with similar shape, a decrease in amplitude prevented the BSS from associating the doublet to the same MU as the main action potential. Nonetheless, an adaptive change in threshold for detection may in the future solve this problem.

Three MUs found by both decomposition procedures had misalignment for discharges >0.5 ms and this influenced the RoA for those MUs. These misalignments are not necessarily errors. The MU action potential train detected at a certain electrode produces time-locked trains in other electrodes that fall in that MU territory, but can also exhibit some jitter from discharge to discharge due to fluctuations in muscle fiber conduction velocity (Stålberg and Sonoo, 1994). In retaining only one firing pattern per MU, we discarded this information on the jitter. Also, one of the three MUs had a satellite potential which showed some size and temporal jitter. The two algorithms may have used either the main potential or the satellite potential as a reference for the alignment, which may then cause misalignments. Note that the results of the automatic decomposition did not undergo any post-processing. Otherwise, some mistakes could have been easily corrected by plotting the firing rate against time to detect any inconsistencies.

Finally, this work validated for the first time BSS decomposition on a very large number of MUs. Previous validation via comparison between surface and intramuscular data was limited to an average of 1 MU per contraction commonly found in the two datasets (Holobar et al., 2010). In this study, rather than two datasets, we compared the decomposition performance when the same signals were independently analysed by two operators using two different procedures. The total number of common MUs was 123, i.e., 31 per electrode array.

### MU population coherence

Our coherence analysis showed that the synaptic input common to the MN pool may have frequency content up to 75 Hz (Fig. 2 B and D) and that the estimated coherence increases with the number of MUs included in the analysis. Therefore, large populations of concurrently active MUs are necessary to infer characteristics of the neural drive. For a certain frequency of the synaptic input to be detected as common (i.e., statistically significant in the coherence plot), the synaptic input has to be sampled at least twice as fast as that frequency component (Lazar and Pnevmatikakis, 2008). Each MN integrates the supraspinal and afferent inputs and discharges an action potential when the net input exceeds the recruitment threshold. Under the assumption of a common input uniformly distributed to the whole MN pool (Farina et al., 2014), the effective sampling frequency of the synaptic input is the cumulative frequency of all active MNs, i.e. the frequency of the spike train obtained pooling all spike trains together. In voluntary sustained contractions, a MN usually discharges less than 40 action potentials per second (Enoka and Fuglevand, 2001). As a result, sampling by few MNs limits the maximal frequency of the signal recorded from the output of the spinal cord, while large populations allow the synaptic input to be reconstructed more accurately from the MN output.

The very large frequency content identified for the neural drive from the spinal cord to muscles is unexpected as muscles can only contract within a narrow bandwidth (<10 Hz) (Baldissera et al., 1998). The issue of the mismatch between the bandwidth of the neural drive and of the muscle dynamics has been previously discussed in relation to the β band (Watanabe and Kohn, 2015). It has long been known that beta oscillations are present in MN output (Ibáñez et al., 2021) while they are filtered out by the muscle contractile properties. The new observation of a much greater frequency content than the β oscillations indicates the variety of common inputs received by the MN pool. Gamma-range cortico-muscular coherence has been observed during strong isometric voluntary contractions (Ushiyama et al., 2012), and during dynamic contractions (Andrykiewicz et al., 2007), suggesting that the gamma-band rhythmic drive from the cortex contributes, at least in part, to the EMG activity at that frequency band. Our results show that human muscles can manifest rhythmic electrical oscillations in the gamma-band also during low intensity isometric contractions.

### Reciprocal influence of motoneuron discharges onto the homonymous pool

Our study included the analysis of the influence of the discharges of early recruited MUs on those recruited later (Fig. 3, R1 → R2) and vice versa (Fig. 3, R2 → R1). We observed that the highest value of the six cross-histograms was obtained at 0 s, indicating the common drive received by the MN pool (De Luca and Erim, 1994). Early recruited MUs were less likely to fire for about 15 ms (Fig. 3A and C, S1 and S3, R2 → R1) or 40 ms (Fig. 3B, S2, R2 → R1) after the discharge of later recruited MUs. This observation fits with recurrent inhibition by Renshaw cells which occurs with similar timing (Bhumbra et al., 2014). Recurrent inhibition has been studied in isolated cells in *in vitro* experiments or in anesthetized animal preparations. The main method to test homonymous recurrent inhibition in humans is indirect and relies on changes in H-reflex modulation caused by presumed recurrent effects (Pierrot-Deseilligny and Burke, 2005). An elegant method to evaluate recurrent inhibition in humans at individual MN level has been proposed by Özyurt et al. (2019). However, this method can only be used to assess the impact of the largest on smaller MUs as it evaluates the effect of electrical stimulation on the background firing of small MUs. On the contrary, our method can be applied in both directions across the MN pool during voluntary contractions. Özyurt et al. (2019) reported an average latency for recurrent inhibition of 37.7 ms from a peripheral stimulus for the soleus muscle, which is compatible with the dips at ∼ 40 ms visible in the cross-histograms of S2 (Fig. 3B). For S1 and S3, inhibition occurred earlier than for S2 (Fig. 3A and C).

In conclusion, we present a novel high-density intramuscular array along with a methodology that fully automatically identifies the spike trains of relatively large number of MUs, unveiling new knowledge behind MN population coding. We demonstrated that the number of automatically identified MUs is high enough to reveal the presence of significant coherence between groups of MNs in the frequency range up to 75 Hz and the effect of Renshaw inhibition on the homonymous MN pool. These results constitute an important step forward in the *in vivo* population coding analysis of the human motor system.

## ACKNOWLEDGEMENTS

The authors would like to thank Marco Beato and Rob Brownstone, University College London, for useful discussion.

## METHODS

### Manufacturing process

The thin-film electrode array structure was built using microfabrication processes. The electrode array was built over a silicon wafer used as a platform for the production. The structure was built layer by layer with layers of metal for tracks sandwiched between three layers of polyimide. Metals were patterned using a photolithography process.

First, a platinum etch mask was deposited and lift-off structured on a 4 inches silicon wafer. In the next step, a 5 μm polyimide layer (PI 2611, HD Microsystems) was spun on the wafer and cured at 350°C. The lower platinum electrode contacts and tracks were then sputtered and lift-off structured. Another 10 μm polyimide layer was deposited, followed by the upper platinum electrode tracks and contacts, which were sputtered and lift-off structured, followed by a final 5 μm polyimide layer for insulation. To reach the contacts on the lower side, the silicon wafer was etched from the backside using reactive ion etching. In a second reactive ion etching step, the lower electrode contacts were opened using the previously deposited platinum layer as etch mask. An aluminum etch mask was then deposited on the top side and used for reactive ion etching of the polyimide to open the contacts on the upper side. After removal of the aluminum mask, the microfabrication process was completed, and the separated double-sided electrode arrays were removed from the wafer using tweezers. The electrode contacts were coated with microrough platinum using electroplating from an aqueous solution of hexachloroplatinic acid (Poppendieck et al., 2014). This reduced the electrode impedance by about one order of magnitude so that the resulting values of impedance spectroscopy were ∼10 kΩ at 1 kHz. A plug (Harwin M50-4902045 connector) was soldered to the adapter as the interface with external hardware. Each electrode array was inserted into a hypodermic needle with the bevel smoothed with a laser (PICCO LASER, O.R. Lasertechnologie, DE).

### Subjects

Three healthy men (age range: 29 - 39 years) participated in the experiment, which was approved by the Ethical Committee of the University Medical Center of Göttingen and conducted according to the Declaration of Helsinki (2008).

### Experimental procedure

The subject was seated in the chair of a Biodex System 3 (Biodex Medical Systems Inc., NY, USA) with the right leg and foot stably fixated. He was asked to perform two brief maximal voluntary contractions with 5 minutes interval in between to recover from fatigue. The peak of the two was considered as the maximal voluntary contraction (MVC). Electrode array placement followed 5 extra minutes of rest. The skin was cleaned with alcohol and the thin-film electrode array(s) were inserted into the middle of the proximal half of the tibialis anterior muscle, perpendicular to the skin with the tip of the needle to a depth of 2.5 cm below the fat layer as estimated by ultrasound (Telemed Ltd. Vilnius, Lithuania). The two electrode arrays in S1 were about 3 and 1 cm distant in the longitudinal and perpendicular direction of the muscle, respectively.

Intramuscular EMG signals were recorded with a multichannel amplifier (EMG-USB2, OT-Bioelettronica, Torino, Italy) with a gain of 200-500, and band-pass filtered (8th order Bessel filter, high-pass cut-off frequency 10-100; low-pass cut-off frequency 4400 Hz), before being sampled at 10 kHz, using a 12-bit A/D converter. The EMG signals were acquired in a unipolar derivation with reference and ground electrodes at the ankle.

The subject was then asked to perform a brief contraction at 20 and 30% MVC during which the experimenters judged the signal quality. Following these trials, S1 was asked to perform a steady contraction at 20% MVC, whereas S2 and S3 were given 30% MVC as the target force level. Subjects were asked to perform a steady contraction lasting at least 1 min. The subject was provided with real-time force feedback displayed on a screen. The target force level was represented as straight line on the computer screen and the force exerted by the subject as a running dot. The subject was instructed to keep the position of the dot as close as possible to the straight line. He was allowed to complete the 1 min contraction at once or in multiple contractions with rest at will in between.

### Signal quality assessment

EMG signals were bandpass-filtered in the bandwidth 100-4400 Hz (third-order Butterworth, zero-lag filter) so that the frequency content was the same for all signals. We quantified the baseline noise as the average across 160 channels (4 electrode arrays x 40 channels / array) of the root-mean-square of a 4 s segment of data recorded at rest.

### Signal decomposition

The recorded signals were independently manually and automatically decomposed into the constituent MU action potential trains by two expert investigators (SM and AH, respectively). In both cases, signals were high pass filtered at 250 Hz prior decomposition. In case of manual decomposition, intramuscular EMG signals from each thin-film array were decomposed using the decomposition software EMGLAB (McGill et al., 2005), that relies on spike sorting to detect MU action potentials. Each channel was decomposed independently and the series of discharges of a single MU were manually edited for resolving missed discharges and superimpositions. This process was conducted for each MU identified from the same channel until the residual signal, obtained by subtracting all averaged MU action potentials from the raw signal, was comparable in power with the raw signal baseline noise, indicating that all MU activity had been accounted for. As the same MU could be detected in adjacent channels, the decomposition results from all channels were then merged by automatically identifying the MUs detected at more than one electrode. Discharge patterns with more than 75% discharges closer than 1 ms were considered to belong to the same MU identified on different channels. Differences in the discharge patterns of the same MU extracted from different channels were examined and resolved by the investigator in charge, so that at the final stage of the manual decomposition, each MU was represented by a unique firing pattern.

A second investigator (AH), automatically decomposed the 20 s signals using the convolution kernel compensation algorithm (CKC) (Holobar and Zazula, 2007). To briefly summarize the algorithm working principle, assuming absence of noise, we can express the intramuscular EMG signal *x*_*c*_*(k)* recorded at channel *c* as the sum of trains of action potentials (one train for each active MU):

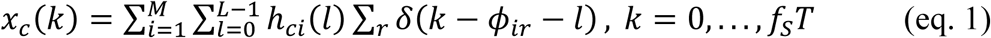

where *f*_*S*_ is the sampling frequency, *T* the signal duration, *h*_*ci*_(*l*) is the action potential of the *i-th* MU as recorded at the *c-th* channel, Σ_*r*_ *δ*(*k* − *ϕ*_*ir*_) the spike train of the *i-th* MU with spikes at times *ϕ*_*ir*_, *L* the duration of the action potentials, and *M* the number of active MUs.

Equation 1 can be re-written in matrix form as follows:

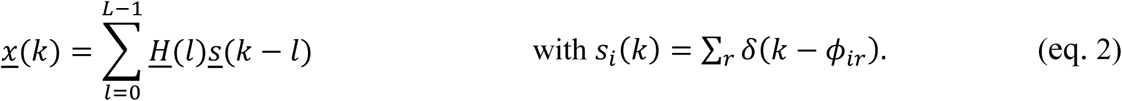

Once the mixing matrix *<u>H</u>* is identified, the source pulse trains can be extracted by multiplying the EMG signals (*<u>x</u>*) by the inverse of *<u>H</u>* (unmixing matrix). The reliability of the automatic decomposition was assessed by the pulse-to-noise ratio, which is a signal-based metric that has been validated to assess the decomposition accuracy of BSS-based decomposition algorithms (Holobar et al., 2014).

### Assessment of the decomposition accuracy

For each electrode array (3 subjects, 4 arrays), we report the number of MUs identified by the manual and automatic decomposition, and those commonly identified by both approaches. We first inspected the results of the manual decomposition. We calculated the RoA (Holobar et al., 2010) between each pair of MU firing patterns identified from the same 40 channel array, to ensure that they were unique. The RoA was defined as the ratio between the number of discharges that were present in both firing patterns (common) and the sum of the number of common discharges and the number of discharges present in only one of the two firing patterns. A tolerance of 10 sample (< 1 ms) was used when identifying common discharges.

Each MU firing pattern was accurately estimated from the comparison between the firing patterns of that MU in multiple channels. To assess the robustness of the estimation, we calculated the multichannel MU action potentials by spike triggered averaging (Farina et al., 2002), i.e., by averaging the EMG of each channel using the discharges obtained from decomposition as a trigger. For each MU, we then counted the number of channels where the peak-to-peak amplitude of the action potential was greater than 10 times the average RMS of the baseline noise across the 40 channels. The higher the number of channels exceeding the threshold, the higher the likelihood that the firing pattern was accurately estimated (Mambrito and De Luca, 1984).

The RoA was also used to check whether there were MUs in common between array 1 and array 2 of S1. As a further check, we performed cross-spike triggered averaging by averaging the EMG of each channel of array1 (array2) using the discharges obtained from decomposition of the EMG from array2 (array1) as a trigger. A temporal support of 20 ms (centered about the MU firing) was used in the spike triggered averaging procedure to account for the propagation delay between the position of the electrode arrays, which were about 3 cm apart. For MUs in common between the two arrays, the cross-averaging procedure will yield an action potential with higher amplitude than the baseline noise.

We then compared the MU firing patterns extracted by the two decomposition procedures (manual and automatic). Here RoA was defined as the ratio between the matched discharges resulting from the comparison of the two procedures and the sum of matched and unmatched discharges. Discharge patterns with more than 75% discharges closer than 0.5 ms were considered to belong to the same MU identified by the manual and automatic procedure (common MU).

### MU population coherence

The discriminated spike trains were used to compute spectral coherence between groups of MUs, with numerosity ranging from 1 to half of the maximum number of extracted MUs. The allocation of MUs into groups was repeated 25 times for each group size (i.e., 1, 2, 3, … MU spike trains) and the average coherence across the 25 repetitions was calculated. For each MU, spike trains were represented with binary vectors of 0 and 1, with 1 indicating the occurrence of a discharge. Within each MU group, the spike trains were summed to provide a cumulative spike train. Coherence analysis was performed on 0.5 s non-overlapping Hanning windows of the cumulative spike trains with a length of the Fast Fourier Transform equal to the sampling rate. To define the significance threshold for coherence peaks, the confidence level CL was calculated as (Rosenberg et al. 1989):

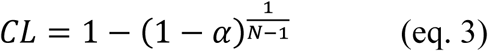

where *N* and α represent the number of segments used in the coherence calculation (data length/number of windows) and the confidence level (95%), respectively.

### Connectivity among motoneurons

Connectivity among MNs was estimated by the cross-histogram of the discharge of pairs of MUs (1 ms resolution). To consider the opposing views on the distribution of recurrent inhibition between early- and late-recruited MUs within the homonymous MN pool (Granit et al., 1957; Haase et al., 1975; Hultborn et al., 1988), we investigated separately higher and lower threshold MUs. MUs were ordered by firing rate based on the fact that at a given force, earlier recruited MUs discharge faster than later recruited ones (De Luca and Erim, 1994). As control conditions, we generated 4 types of firing patterns with the same number of discharges as the detected MUs in the same time interval and *i)* uniformly distributed discharge times, *ii)* equal inter-spike intervals, *iii)* discharged times obtained from the experimental ones by applying a time shift of 0 to 70 ms to the whole MU action potential train (different for the different MUs, but the same for all action potentials of the same MU), and *iv)* discharged times obtained from the experimental ones by adding or subtracting a time in the range of 0 to 10% of the average inter-spike interval for each MU (different time shifts for each individual action potential).

## Notes

### Competing Interest Statement

The authors have declared no competing interest.

## REFERENCES

Adrian, E.D., and Bronk, D.W. (1929). The discharge of impulses in motor nerve fibres. Part II. The frequency of discharge in reflex and voluntary contractions. J. Physiol. 67, 119–151.

Alvarez, F.J. (2019). A motor physiology recurrent topic: simplify assumptions to gain extra insight. J. Physiol. 597, 2117–2118.

Andrykiewicz, A., Patino, L., Naranjo, J.R., Witte, M., Hepp-Reymond, M.C., and Kristeva, R. (2007). Corticomuscular synchronization with small and large dynamic force output. BMC Neurosci. 8, 1–12.

Baldissera, F., Cavallari, P., and Cerri, G. (1998). Motoneuronal pre-compensation for the low-pass filter characteristics of muscle. A quantitative appraisal in cat muscle units. J. Physiol. 511, 611–627.

Basmajian, J., and Stecko, G. (1962). A new bipolar electrode for electromyography. J. Appl. Physiol. 17, 849.

Bhumbra, G.S., Bannatyne, B.A., Watanabe, M., Todd, A.J., Maxwell, D.J., and Beato, M. (2014). The recurrent case for the Renshaw cell. J. Neurosci. 34, 12919–12932.

Connelly, D.M., Rice, C.L., Roos, M.R., and Vandervoort, A.A. (1999). Motor unit firing rates and contractile properties in tibialis anterior of young and old men. J. Appl. Physiol. 87, 843–852.

Denslow, J.S. (1948). Double discharges in human motor units. J. Neurophysiol. 11, 209–215.

Desmedt, J. (1973). New developments in electromyography and clinical neurophysiology (Basel: S Karger AG).

Eccles, J.C., Eccles, R.M., Iggo, A., and Lundberg, A. (1961). Electrophysiological investigation of Renshaw cells. Electrophysiol. Investig. Renshaw Cells 159, 461–478.

Enoka, R.M. (1995). Morphological features and activation patterns of motor units. J. Clin. Neurophysiol. 12, 538–559.

Enoka, R.M. (2019). Physiological validation of the decomposition of surface EMG signals. J. Electromyogr. Kinesiol. 46, 70–83.

Enoka, R.M., and Fuglevand, A.J. (2001). Motor unit physiology: some unresolved issues. Muscle Nerve 24, 4–17.

Erim, Z., De Luca, C.J., Mineo, K., and Aoki, T. (1996). Rank-ordered regulation of motor units. Muscle and Nerve 19, 563–573.

Farina, D., Arendt-Nielsen, L., Merletti, R., and Graven-Nielsen, T. (2002). Assessment of single motor unit conduction velocity during sustained contractions of the tibialis anterior muscle with advanced spike triggered averaging. J. Neurosci. Methods 115, 1–12.

Farina, D., Yoshida, K., Stieglitz, T., and Koch, K.P. (2008). Multichannel thin-film electrode for intramuscular electromyographic recordings. J. Appl. Physiol. 104, 821–827.

Farina, D., Holobar, A., Merletti, R., and Enoka, R.M. (2010). Decoding the neural drive to muscles from the surface electromyogram. Clin. Neurophysiol. 121, 1616–1623.

Farina, D., Negro, F., and Dideriksen, J.L. (2014). The effective neural drive to muscles is the common synaptic input to motor neurons. J. Physiol. 592, 3427–3441.

Granit, R., Pascoe, J.E., and Steg, G. (1957). The behaviour of tonic α and β motoneurones during stimulation of recurrent collaterals. J. Physiol. 138, 381–400.

Haase, J., Cleveland, S., and Ross, H.G. (1975). Problems of postsynaptic autogenous and recurrent inhibition in the mammalian spinal cord. Rev. Physiol. Biochem. Pharmacol. 73, 73–129.

Hassler, C., Boretius, T., and Stieglitz, T. (2011). Polymers for neural implants. J. Polym. Sci. Part B Polym. Phys. 49, 18–33.

Holobar, A., and Zazula, D. (2007). Multichannel blind source separation using convolution kernel compensation. IEEE Trans. Signal Process. 55, 4487–4496.

Holobar, A., Farina, D., Gazzoni, M., Merletti, R., and Zazula, D. (2009). Estimating motor unit discharge patterns from high-density surface electromyogram. Clin. Neurophysiol. 120, 551–562.

Holobar, A., Minetto, M.A., Botter, A., Negro, F., and Farina, D. (2010). Experimental analysis of accuracy in the identification of motor unit spike trains. IEEE Trans. Neural Syst. Rehabil. Eng. 18, 221–229.

Holobar, A., Minetto, M.A., and Farina, D. (2014). Accurate identification of motor unit discharge patterns from high-density surface EMG and validation with a novel signal-based performance metric. J. Neural Eng. 11, 016008.

Hultborn, H., and Pierrot-Deseilligny, E. (1979). Changes in recurrent inhibition during voluntary soleus contractions in man studied by an H-reflex technique. J. Physiol. 297, 229–251.

Hultborn, H., Lindström, S., and Wigström, H. (1979). On the function of recurrent inhibition in the spinal cord. Exp. Brain Res. 37, 399–403.

Hultborn, H., Katz, R., and Mackel, R. (1988). Distribution of recurrent inhibition within a motor nucleus. II. Amount of recurrent inhibition in motoneurones to fast and slow units. Acta Physiol. Scand. 134, 363–374.

Ibáñez, J., Del Vecchio, A., Rothwell, J.C., Baker, S.N., and Farina, D. (2021). Only the fastest corticospinal fibers contribute to β corticomuscular coherence. J. Neurosci. 41, 4867–4879.

Lazar, A.A., and Pnevmatikakis, E.A. (2008). Faithful representation of stimuli with a population of integrate-and-fire neurons. Neural Comput. 20, 2715–2744.

Lazar, A.A., and Tóth, L.T. (2004). Perfect recovery and sensitivity analysis of time encoded bandlimited signals. IEEE Trans. Circuits Syst. I Regul. Pap. 51, 2060–2073.

LeFever, R.S., and De Luca, C.J. (1982). A procedure for decomposing the myoelectric signal into its constituent action potentials - Part I: technique, theory, and implementation. IEEE Trans. Biomed. Eng. BME-29, 149–157.

De Luca, C.J., and Erim, Z. (1994). Common drive of motor units in regulation of muscle force. Trends Neurosci. 17, 299–305.

Mambrito, B., and De Luca, C.J. (1984). A technique for the detection, decomposition and analysis of the EMG signal. Electroencephalogr. Clin. Neurophysiol. 58, 175–188.

McGill, K.C., Lateva, Z.C., and Marateb, H.R. (2005). EMGLAB: An interactive EMG decomposition program. J. Neurosci. Methods 149, 121–133.

Muceli, S., Poppendieck, W., Negro, F., Yoshida, K., Hoffmann, K.P., Butler, J.E., Gandevia, S.C., and Farina, D. (2015). Accurate and representative decoding of the neural drive to muscles in humans with multi-channel intramuscular thin-film electrodes. J. Physiol. 593, 3789–3804.

Muceli, S., Poppendieck, W., Hoffmann, K.P., Dosen, S., Benito-León, J., Barroso, F.O., Pons, J.L., and Farina, D. (2019). A thin-film multichannel electrode for muscle recording and stimulation in neuroprosthetics applications. J. Neural Eng. 16, 026035.

Negro, F., Muceli, S., Castronovo, A.M., Holobar, A., and Farina, D. (2016). Multi-channel intramuscular and surface EMG decomposition by convolutive blind source separation. J. Neural Eng. 13, 026027.

Özyurt, M.G., Piotrkiewicz, M., Topkara, B., Weisskircher, H.W., and Türker, K.S. (2019). Motor units as tools to evaluate profile of human Renshaw inhibition. J. Physiol. 597, 2185–2199.

Pierrot-Deseilligny, E., and Burke, D. (2005). Recurrent inhibition. In The Circuitry of the Human Spinal Cord, (New York: Cambridge University Press), pp. 151–196.

Poppendieck, W., Sossalla, A., Krob, M.O., Welsch, C., Nguyen, T.A.K., Gong, W., DiGiovanna, J., Micera, S., Merfeld, D.M., and Hoffmann, K.P. (2014). Development, manufacturing and application of double-sided flexible implantable microelectrodes. Biomed. Microdevices 16, 837–850.

Poppendieck, W., Muceli, S., Dideriksen, J., Rocon, E., Pons, J.L., Farina, D., and Hoffmann, K.P. (2015). A new generation of double-sided intramuscular electrodes for multi-channel recording and stimulation. Proc. Annu. Int. Conf. IEEE Eng. Med. Biol. Soc. EMBS 2015-Novem, 7135–7138.

Renshaw, B. (1941). Influence of discharge of motoneurons upon excitation of neighboring motoneurons. J. Neurophysiol. 4, 167–183.

Stålberg, E. V., and Sonoo, M. (1994). Assessment of variability in the shape of the motor unit action potential, the “jiggle,” at consecutive discharges. Muscle Nerve 17, 1135–1144.

Stieglitz, T., Beutel, H., Schuettler, M., and Meyer, J.-U. (2000). Micromachined, polyimide-based devices for flexible neural interfaces. Biomed. Microdevices 2, 283–294.

Ushiyama, J., Masakado, Y., Fujiwara, T., Tsuji, T., Hase, K., Kimura, A., Liu, M., and Ushiba, J. (2012). Contraction level-related modulation of corticomuscular coherence differs between the tibialis anterior and soleus muscles in humans. J. Appl. Physiol. 112, 1258–1267.

Del Vecchio, A., Holobar, A., Falla, D., Felici, F., Enoka, R.M., and Farina, D. (2020). Tutorial: Analysis of motor unit discharge characteristics from high-density surface EMG signals. J. Electromyogr. Kinesiol. 53, 102426.

Watanabe, R.N., and Kohn, A.F. (2015). Fast oscillatory commands from the motor cortex can be decoded by the spinal cord for force control. J Neurosci 35, 13687–13697.

